# Identification of Pathways Associated with Chemosensitivity through Network Embedding

**DOI:** 10.1101/168450

**Authors:** Sheng Wang, Edward Huang, Junmei Cairns, Jian Peng, Liewei Wang, Saurabh Sinha

## Abstract

Basal gene expression levels have been shown to be predictive of cellular response to cytotoxic treatments. However, such analyses do not fully reveal complex genotype-phenotype relationships, which are partly encoded in highly interconnected molecular networks. Biological pathways provide a complementary way of understanding drug response variation among individuals. In this study, we integrate chemosensitivity data from a recent pharmacogenomics study with basal gene expression data from the CCLE project and prior knowledge of molecular networks to identify specific pathways mediating chemical response. We first develop a computational method called PACER, which ranks pathways for enrichment in a given set of genes using a novel network embedding method. It examines known relationships among genes as encoded in a molecular network along with gene memberships of all pathways to determine a vector representation of each gene and pathway in the same low-dimensional vector space. The relevance of a pathway to the given gene set is then captured by the similarity between the pathway vector and gene vectors. To apply this approach to chemosensitivity data, we identify genes with basal expression levels in a panel of cell lines that are correlated with cytotoxic response to a compound, and then rank pathways for relevance to these response-correlated genes using PACER. Extensive evaluation of this approach on benchmarks constructed from databases of compound target genes, compound chemical structure, as well as large collections of drug response signatures demonstrates its advantages in identifying compound-pathway associations, compared to existing statistical methods of pathway enrichment analysis. The associations identified by PACER can serve as testable hypotheses about chemosensitivity pathways and help further study the mechanism of action of specific cytotoxic drugs. More broadly, PACER represents a novel technique of identifying enriched properties of any gene set of interest while also taking into account networks of known gene-gene relationships and interactions.

## INTRODUCTION

Large-scale cancer genomics projects, such as the Cancer Genome Atlas^1^, the Cancer Genome project^2^, and the Cancer Cell Line Encyclopedia project^3^, and cancer pharmacology projects such as the Genomics of Drug Sensitivity in Cancer project^4^ have generated a large volume of genomics and pharmacological profiling data. As a result, there is an unprecedented opportunity to link pharmacological and genomic data to identify therapeutic biomarkers^5–7^. In pursuit of this vision, significant efforts have been invested in identifying the genetic basis of drug response variation among individual patients. For instance, a recent study performed a comprehensive survey of genes with basal expression levels in cancer cell lines that correlate with drug sensitivity, revealing potential gene candidates for explaining mechanisms of action of various drugs^8^.

While significant efforts have focused on specific genes that interact with compounds and confer observed cellular phenotypes, there has been relatively little progress in studying the synergistic effects of genes. These effects are key factors in comprehensively deciphering the mechanisms of action of compounds and understanding complex phenotypes^9^. Similarly, pathways, which comprise a set of interacting genes, have emerged as a useful construct for gaining insights into cellular responses to compounds. Analysis at the pathway level not only reduces the analytic complexity from tens of thousands of genes to just hundreds of pathways, but also contains more explanatory power than a simple list of differentially expressed genes^10^. Consequently, an important yet unsolved problem is the effective identification of pathways mediating drug response variation. Although the associated pathways for certain drugs have been studied experimentally^11–13^, *in vitro* pathway analysis is costly and inherently difficult, making it hard to scale to hundreds of compounds.

Fortunately, a growing compendium of genomic, proteomic, and pharmacologic data allows us to develop scalable computational approaches to help solve this problem. Although statistical significance tests and enrichment analyses can be naturally applied to compound-pathway association identification (e.g., by testing the overlap between pathway members and differentially expressed genes), these approaches fail to leverage well-established biological relationships among genes^14–17^. Even when analyzing individual genes, molecular networks such as protein-protein interaction networks have been shown to play crucial roles in understanding the cellular drug response^9, 18–21^. Therefore, we propose to combine similar molecular networks with gene expression and drug response data for pathway identification. However, integrating these heterogeneous data sources is statistically challenging. Moreover, networks are high-dimensional, incomplete, and noisy. Thus, our algorithm needs to accurately and comprehensively identify pathway while exploiting suboptimal networks.

In this work, we present PACER, a novel, network-assisted algorithm that identifies pathway associations for any gene set of interest. Additionally, we apply the algorithm to discover chemosensitivity-related pathways. PACER first constructs a heterogeneous network that includes pathways and genes, pathway membership information, and gene-gene relationships from a molecular network such as protein-protein interaction network. It then applies a novel dimensionality reduction algorithm to this heterogeneous network to obtain compact, low-dimensional vectors for pathways and genes in the network. Pathways that are topologically close to drug response-related genes in the network are co-localized with those genes in this low-dimensional vector space. Hence, PACER ranks each pathway based on its proximity in the low-dimensional space to genes that have basal expressions highly correlated with drug response. We evaluated PACER's ability to identify compound-pathway associations with three ‘ground truth’ sets built from compound target data^8^, compound structure data^22^, and LINCS differential expression data^23^. When comparing PACER to state-of-the-art methods that ignore prior knowledge of interactions among genes, we observed substantial improvement of the concordance with the chosen benchmarks. Even though we developed PACER and tested its ability to identify compound-pathway associations, the algorithm is applicable to any scenario in which one seeks to discover pathways related to a pre-specified gene set of interest, while utilizing a given gene network.

## MATERIALS AND METHODS

### Compound response data and gene expression data

We obtained a large-scale compound response screening dataset from Rees *et al.*^8^, which spans 481 chemical compounds and 842 human cancer cell lines encompassing 25 lineages. These 481 compounds were collected from different sources including clinical candidates, FDA-approved drugs and previous chemosensitivity profiling experiments. Area under the drug response curve (AUC) was used by the authors of that study to measure cellular response to individual compound. We also obtained gene expression profiles for these cell lines from the Cancer Cell Line Encyclopedia (CCLE) project^3^, profiled using the GeneChip Human Genome U133 Plus 2.0 Array. Since these expression measurements were done in each cell line without any drug treatment, they are referred to as ‘basal’ expression levels. In contrast, the expression profiling of a cell line was performed *after* treatment with a drug in certain studies^23^. We obtained the SMILE specification of each drug from PubChem^22^ and then calculated the Tanimoto similarity scores between all pairs of drugs based on their SMILE specifications.

### STRING-based molecular network and NCI pathway collection

We obtained a collection of six human molecular networks from the STRING database v9.1^24^. These six networks include experimentally derived protein-protein interactions, manually curated protein-protein interactions, protein-protein interactions transferred from model organism based on orthology, and interactions cmputed from genomic features such as fusion-fusion events, functional similarity and co-expression data. There are 16,662 genes in the network. We used all the STRING channels except “text-mining” and used the Bayesian integration method provided by STRING. Since our approach can deal with different edge weights, we did not set a threshold to remove low confidence edges. We referred to this integrated network as the ‘STRING-based molecular network’. To test whether genes that are highly correlated with many compounds tend to have higher degrees in the network, we formed two groups of genes. One group contained genes that are correlated with over 100 compounds, and the other group contained the remaining genes. We then used the Wilcoxon signed-rank test to test whether the degrees of genes in these two groups were from the same distribution. We obtained a collection of 223 cancer-related pathways from the National Cancer Institute (NCI) pathway database. These manually curated pathways include human signaling and regulatory pathways as well as key cellular processes^25^.

### The PACER Framework

PACER integrates pathway information with the STRING-based molecular network described above by constructing a heterogeneous network of genes and pathways. An edge exists between two genes if they are connected in the network. An edge exists between a pathway and a gene if the gene belongs to the pathway. There are no direct pathway-pathway edges in the heterogeneous network. PACER adopts diffusion component analysis (DCA), a recently developed network representation algorithm to learn a low-dimensional vector for each node in the network^26^. Because of its ability to handle noisy and missing edges in the biological network, DCA has achieved state-of-the-art results in different tasks^26, 27^. Since compounds are not nodes in the constructed heterogeneous network, only genes and pathways are projected onto the low-dimensional space. After learning the low-dimensional representations of all nodes (genes and pathways), DCA ranks pathways based on the weighted cosine similarities between a pathway and the set of 250 genes most correlated with response to a compound. We henceforth refer to this set of genes as “response-correlated genes” (RCG) for the compound. These genes’ expression values are most significantly correlated with chemosensitivity. We found that the performance of the PACER method is stable for different choices (200, 250, and 300) for the number of RCGs considered in this step (**Suppl. Figure 7-9.**).

### LINCS drug perturbation profiles

LINCS is a data repository of over 1.3 million genome-wide expression profiles of human cell lines subjected to a variety of perturbation conditions, which include treatments with more than 20 thousand unique compounds at various concentrations. Each perturbation experiment is represented by a list of differentially expressed genes that are ranked based on *z*-scores of perturbation expression relative to basal expression. For each gene, we first took the difference between its expression in a perturbation condition and its expression in a control condition (i.e., treatment with pure DMSO solvent). We then considered the differential expression of the gene in multiple perturbation experiments involving that compound (i.e., different concentrations, time points, and cell lines). We used the maximum differential expression to represent the compound's effect on that gene's expression. All genes were then ranked by their differential expression on treatment with the compound, and the top 250 genes were treated as differentially expressed genes (DEGs) of the compound, provided their *z*-score has an absolute value greater than 2.

### Comparison with method of Huang *et al.*

We implemented the method of Huang *et al.*^16^ ourselves using the exact same input (i.e., chemosensitivity and gene expression data) as PACER. We first computed a gene's correlation to a drug by calculating the Pearson correlation coefficient between the gene's expression values and the drug response values across cell lines. Let the set of genes in pathway *p* be denoted by *G_p_*, and their correlation values to a drug *d* by *C*(*G_p_*, *d*). Conversely, the set of genes not in pathway *p* is denoted as 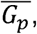 and their correlation values to *d* as *c*(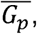 *d*). We then performed the Kruskal-Wallis H-test, following Huang *et al.*, to test if the medians of *C*(*G_p_*, *d*) and *C*(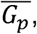 *d*) were significantly different. We used the resulting *p*-value to rank pathways for each drug.

### Software availability

The PACER software is currently available at https://github.com/KnowEnG/PACER. It will also be available through the cloud-based analysis framework KnowEnG (knoweng.org) upon publication.

## RESULTS

### Global analysis of correlations between basal gene expression and compound response

Following the work of Rees *et al.*^8^, we first examined correlations between the compound sensitivity and basal gene expression profiles across hundreds of cell lines. We calculated Pearson correlation coefficients between each gene's expression and the cellular response (measured as the area under the curve or AUC) to each compound, across different cell lines (**Figure 1A**). Compared to the IC50 and EC50 scores, AUC simultaneously captures the efficacy and potency of a drug. Of the ~8.7 million pairs of genes and compounds tested, we found 294,789 to be significantly correlated (*p*-value < 0.0001 after Bonferroni correction, corresponding to a Pearson correlation coefficient of 0.215). Within these significantly correlated pairs, 1,749 genes were correlated with over 100 compounds (**Figure 1B, Suppl. Table 1**). We note that these key genes tend to be high-degree nodes in STRING-based molecular network (Wilcoxon rank sum test *p*-value < 9.6e-14, see Methods). We also found that some (10 of 481) compounds were significantly correlated (Pearson correlation *p*-value < 0.0001 after Bonferroni correction) with more than 3,200 genes (**Figure 1C**). Five of these ten compounds are chemotherapeutic agents (**Suppl. Table 2**). In contrast, about 100 compounds were not significantly correlated with any genes; these compounds are mostly probes that either lack FDA approval or are not clinically used. The large disparity among the examined compounds in terms of the number of correlated genes reflects the diversity of these 481 small molecules. While many of them are chemotherapeutic, which can affect the expression of a large number of genes, some compounds may be targeting specific mutations, post-translational modifications, or protein expression. A closer examination revealed that the compounds with the highest cytotoxicity had the fewest gene correlations (i.e., fewest genes whose expression correlates with cytotoxic response were mostly those with low cytotoxicity) (**Suppl. Figure 1**). This suggests that the strategy of identifying compound-associated genes by correlating basal gene expression profiles with cytotoxicity is likely to be more effective for more potent compounds, for which average response is stronger. Note that the gene expression profiles used here are basal and not in response to treatment with compound, hence it was not clear *a priori* that more effective compounds would have larger numbers of gene correlates. In summary, examination of individual genes’ correlations with chemical response confirmed previous reports^4, 8, 28^ that basal gene expression significantly correlates with cytotoxicity across cell lines, especially for effective cytotoxic drugs.

**Figure 1.**
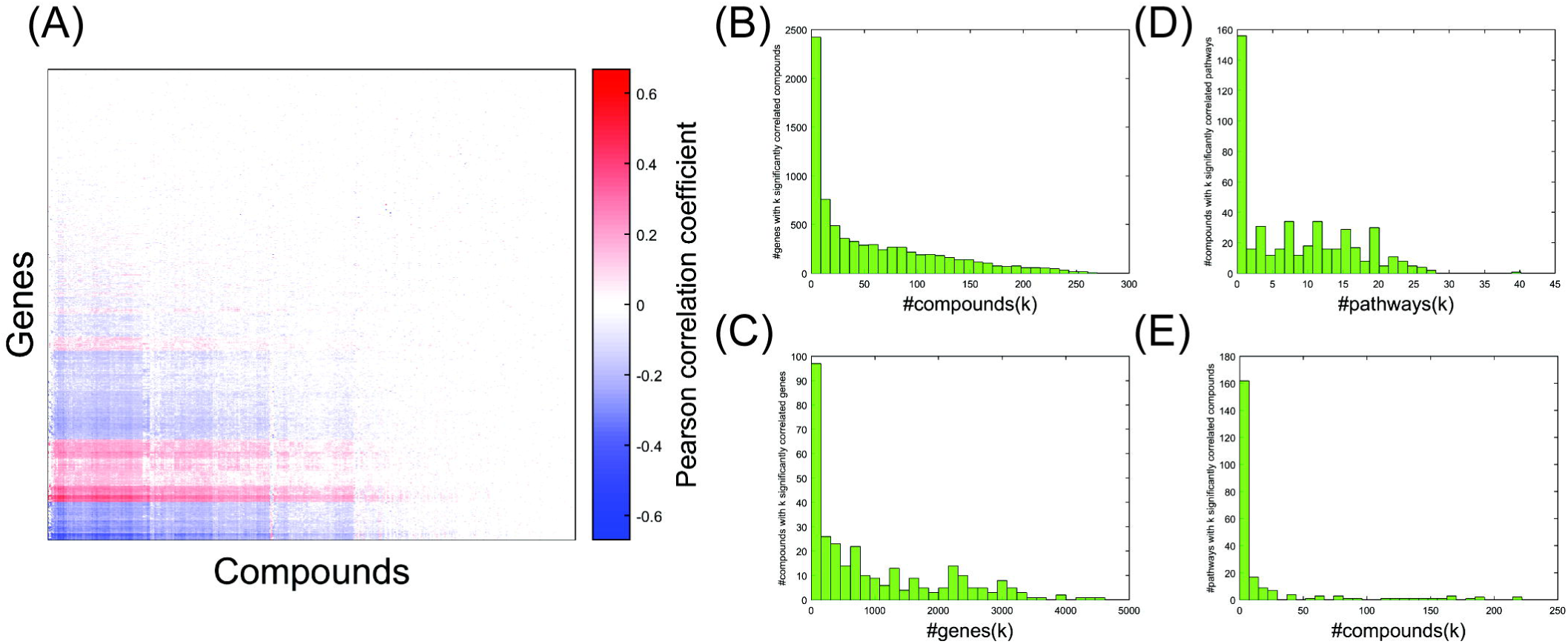
Global analysis of correlations between basal gene expression and compound response. (A) Heatmap of the Pearson correlation coefficient between genes (expression) and compounds (chemosensitivity, measured by AUC values). (B) Histogram of the number of compounds associated with each gene. The *y*-axis shows the number of genes associated with *k* compounds, where *k* is shown on the *x*-axis. (C) Histogram of the number of genes associated with each compound. The *y*-axis shows the number of compounds associated with *k* genes, where *k* is shown on the *x*-axis. (D) Histogram of the number of compounds significantly associated with each pathway (Fisher's exact test FDR ≤ 0.05). (E) Histogram of the number of pathways significantly associated with each compound (Fisher's exact test FDR ≤ 0.05).

### Identifying compound-specific pathways via enrichment tests

The above evidence for correlations between basal gene expression and chemical response raised the possibility that one might discover important biological pathways associated with the response by a systems-level analysis of gene expression data. To explore this, we considered a collection of 223 cancer-related pathways from the National Cancer Institute (NCI) pathway database^25^ and used Fisher's exact test to quantify the overlap between the set of genes in a pathway *p* and (RCGs). A significantly large overlap between the two sets indicates an association between the pathway and the compound. We performed a multiple hypothesis correction on all pathway association tests for each compound, using FDR ≤ 0.05. The results of this baseline method for predicting pathway associations are shown in **Figure 1D** (distribution of the number of compounds that are significantly associated with each pathway) and **Figure 1E** (distribution of the number of pathways significantly associated with each compound). Both distributions revealed a long tail. For instance, while each pathway was associated with an average of 18 compounds (of the 481 tested), there were 10 pathways that were associated with over 150 compounds (**Suppl. Table 3**). Likewise, while each compound was associated with an average of eight pathways, there were 12 compounds associated with over 25 pathways (**Suppl. Table 4**). We show the details of these long tails in **Suppl. Figure 2**.

### A new method for identifying pathways associated with chemical response, based on network embedding

We observed above that key RCGs – those correlated with many compounds – tend to be enriched in high degree nodes of the STRING-based molecular network. This suggests that an analysis combining this network with pathway enrichment tests might provide additional insights. We therefore developed a novel network-based method, called PACER, for scoring compound-pathway associations. PACER (**Figure 2A**) first constructs a heterogeneous network consisting of genes and pathways as nodes. In this network, gene-pathway edges denote pathway memberships based on a compendium of pathways and gene-gene edges from the STRING-based molecular network introduced above (also see Methods). PACER then creates a low-dimensional vector representation for each gene and pathway node in the heterogeneous network, reflecting the node's position in this heterogeneous network. This is done by the Diffusion Component Analysis (DCA) approach reported in previous work^26, 27^. Nodes (i.e., pathways or genes) will have similar vector representations if they are near each other in the network. For instance, two pathway nodes will have similar vector representations if the pathways share genes and/or their genes are related by the STRING-based molecular network. In a similar vein, two genes will have similar representations if they belong to the same pathway(s) and/or exhibit the same network neighbors. A gene and a pathway can also be compared in the low-dimensional space, and will be deemed similar if the gene is in the pathway and/or the gene is related by network to other genes of the pathway. DCA performs network-based embedding without utilizing gene expression or chemical response data. Next, PACER identifies RCGs as a fixed number of genes the expression of which shows the greatest correlation with chemical response to a specific compound. Finally, it scores a pathway based on the average cosine similarity between the vector representation of the pathway and those of the RCGs. A pathway can thus be found to be associated with a compound if, in the network, the pathway genes are closely related to the compound's RCGs; this association can be discovered even if the pathway does not actually include the RCGs. We note that scores assigned by PACER are not statistical significance scores and are meant only to rank pathways for association with a given compound. Also, a negative score assigned to a compound-pathway pair does not imply a negative correlation between expression levels of pathway genes and chemosensitivity. Rather, it only implies a lack of evidence for an association between the compound-pathway pair.

**Figure 2.**
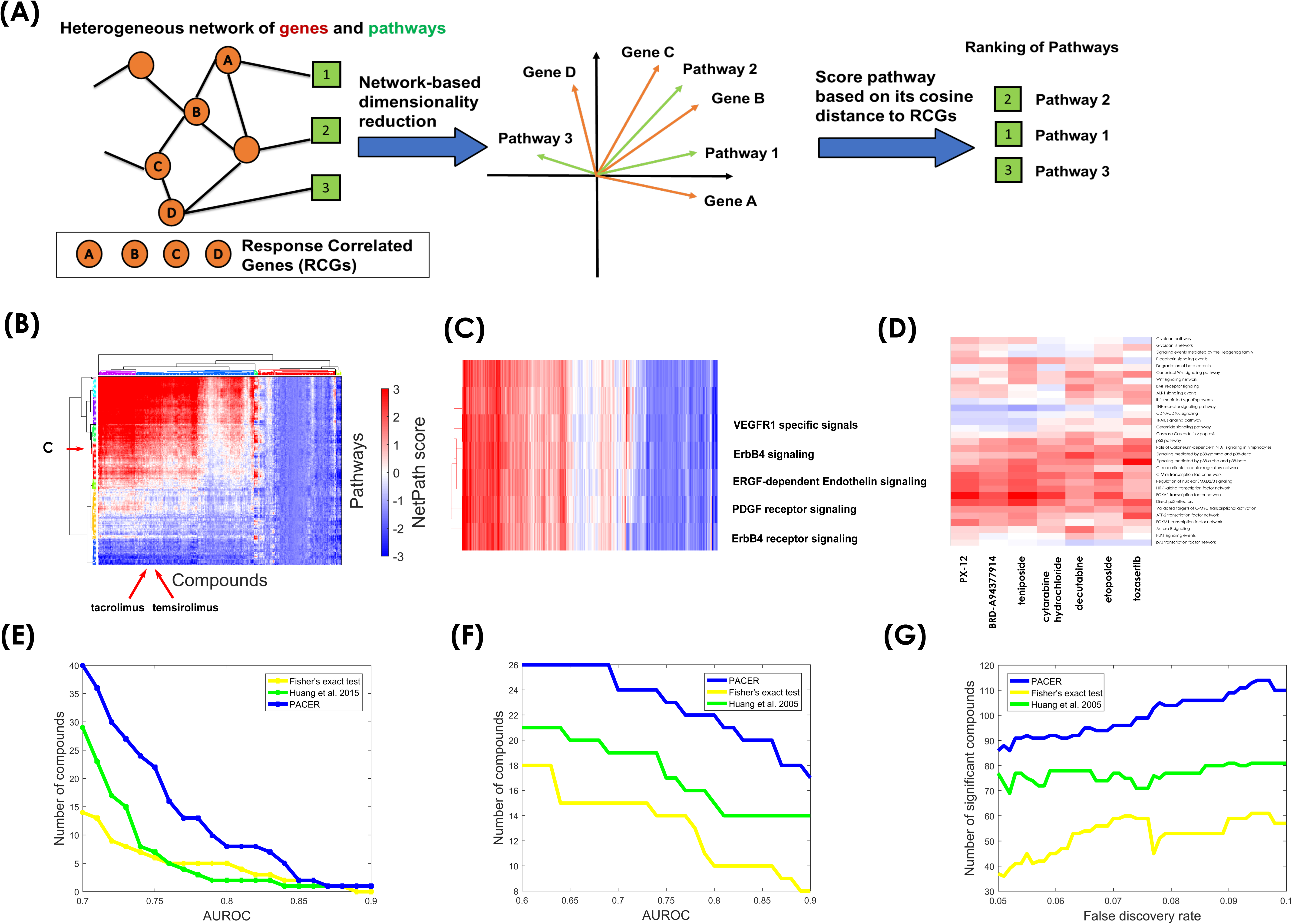
Identifying pathways associated with chemical response using PACER. (A) Schematic description of PACER. (B) Heat map of associations between compounds and pathways (PACER scores). Rows are pathways and columns are compounds. (C) Detailed view of a subset of the red branch cluster of pathways, marked by ‘C’ in Figure 2B. (D) Detailed view of a subnet of purple branch cluster of compounds, marked by a rectangle in Figure 2B. (E) Comparative evaluation of different methods for predicting compound-pathway associations. The ground truth used here is the pathways that contain any known target gene of the compound. (F) Comparison of PACER, Fisher's exact test and Huang *et al.* on predicting compounds with similar chemical structure. The *y*-axis shows the number of compounds with an AUROC larger than *k*, where *k* is shown on the *x*-axis. Compound structure similarity is determined by the Tanimoto similarity score calculated based on their SMILE specifications. Prediction is made according to the Spearman correlation between the pathway rankings of two compounds. (G) Number of compounds with significant overlap (*p* < 0.05) between pathways from LINCS and pathways from PACER, from Huang *et al.* 2005 and from the baseline method (Fisher's exact test) respectively, at different levels of stringency in pathway prediction. Stringency refers to the FDR control used by the baseline method in determining significant pathways. Both PACER and the Huang *et al.* 2005 method were used to predict the same number of (highest scoring) pathways as the baseline method, for a fair comparison.

The PACER association scores for all combinations of 481 compounds and 223 NCI signaling pathways are shown in **Figure 2B**. The pathways cluster into many distinct groups, each with different compound association profiles. One group (cyan branches in the row dendrogram), associated with more than half of the compounds, consists of pathways describing various integrin cell surface interactions (e.g., ‘Integrins in angiogenesis’ pathway, ‘Alpha 4 Beta1 integrin cell surface interactions’ pathway). These pathways are known to play crucial roles in communications among cells in response to small molecules^29^. Notably, integrins are a major family of cell surface adhesion receptors, and are involved in major pathways that contribute to cancer cell survival and resistance to chemotherapy^30^. PACER found 329 of the 481 compounds to be associated with the “integrin family cell surface interactions” pathway. Since PACER scores are not easily assigned statistical significance levels, we chose for each compound *n* pathways with the highest PACER scores, where *n* is the number of statistically significant pathway associations (FDR ≤ 0.05) found by the baseline method above for the same compound. We found literature support for some of these associations. For example, ruxolitinib, a JAK/STAT inhibitor, is associated with integrin pathways by PACER analysis. In a previous study, it was shown that beta 4 integrin enhances activation of the transcription factor STAT3, which is a target of ruxolitinib^31^. Furthermore, vorinostat, a member of a larger class of compounds that inhibit histone deacetylases (HDAC), can induce integrin α5β1 expression and activate MET, leading to resistance^32^. **Figure 2C** reveals another example of functionally related pathways being grouped together. The pathways ‘VEGFR1 specific signals’, ‘ErbB4 signaling events’, ‘EGFR-dependent Endothelin signaling events’, ‘ErbB receptor signaling network’, and ‘PDGF receptor signaling network’ form one group (red branches in row dendrogram), and are associated with masitinib (PDGFRB inhibitor) and RAF265 (VEGFR2 inhibitor), among other compounds (see **Suppl. Table 5**). Vascular endothelial growth factor receptor (VEGFR inhibitor) and platelet-derived growth factor receptor (PDGFR inhibitor) are both members of the family of 58 known tyrosine kinase receptors in humans^33^. Tyrosine kinases have various modulatory functions in growth factor signaling and several of their inhibitors are known for their anti-tumor activity^34^.

**Figure 2B** also shows compounds clustered into different groups based on their associations with pathways. We found that many compounds with similar structure were grouped together. For example, teniposide and etoposide had a Tanimoto similarity score of 0.94 between their SMILE specifications, which was substantially higher than the average Tanimoto similarity score of 0.3716 for all pairs of drugs. They were clustered together in the same group (**Figure 2D**, also marked as a rectangle in **Figure 2B**), which had seven compounds. This group is associated with a set of similar pathways, including ‘p53 pathway’, ‘Direct p53 effectors’, ‘Signaling mediated by p38-alpha and p38-beta’, and ‘Signaling mediated by p38-gamma and p38-delta’. We found support in the literature in favor of some of these associations. For example, a previous study reported that etoposide activates p38MAPK and can be used as a new combined treatment approach when used with p38MAPK inhibitor SB203580^35^. To take another example, temsirolimus and tacrolimus, which are both epipodophyllotoxins and inhibit topoisomerase II, have a Tanimoto similarity score of 0.82, and are grouped closely in **Figure 2B**.

### PACER improves pathway identification

We noted a substantial degree of complementarity between the top predictions of PACER and those of the baseline method that uses Fisher's exact test between RCGs and pathway genes (see **Suppl. Table 5**). For instance, PACER found that PD153035, an ErbB2 inhibitor, is associated with the ‘C-MYC pathway’, reflecting the fact that PD153035 is able to reduce c-Myc protein levels in breast tumor cells^36^. The baseline approach did not find this association to be significant. Similarly, PACER reported that the ‘EGFR-dependent Endothelin signaling events’ pathway is associated with EGFR inhibitor gefitinib^37^, while the baseline method did not.

For a more systematic comparison between the two methods, we evaluated PACER based on a database of known compound targets. We performed the evaluation under the assumption that a pathway containing at least one known target is an associated pathway. Huang *et al.* used and suggested this approach^16^. We used it here to evaluate PACER, the baseline method, as well as a third method presented by Huang *et al.*^16^ Although this third method was proposed to detect association between pathways and drug clades, it can directly detect pathway-compound associations. We implemented the method ourselves (see Methods) and included it in our evaluations. We obtained the known targets for 246 compounds in our compound set from Rees *et al.*^8^ We then computed the AUROC of pathway predictions made by PACER for each compound, and plotted this information alongside analogous information for the baseline method and the method of Huang *et al.*^16^ As shown in **Figure 2E**, PACER identified pathways with higher AUROC compared to the other two methods. For example, PACER identified pathways with an AUROC greater than 0.75 for 22 different compounds, while the baseline method achieved this level of AUROC for only 7 compounds. **Table 1** shows the 10 compounds for which PACER achieved highest AUROC. We present a closer examination of PACER predictions for two of these compounds: ‘cyclophosphamide’ (0.80 AUROC) and ‘nsc23766’ (0.84 AUROC) in **Table 2** and **Table 3**, respectively. These tables also show the complementarity between PACER predictions and those of the baseline method.

**Table 1.**
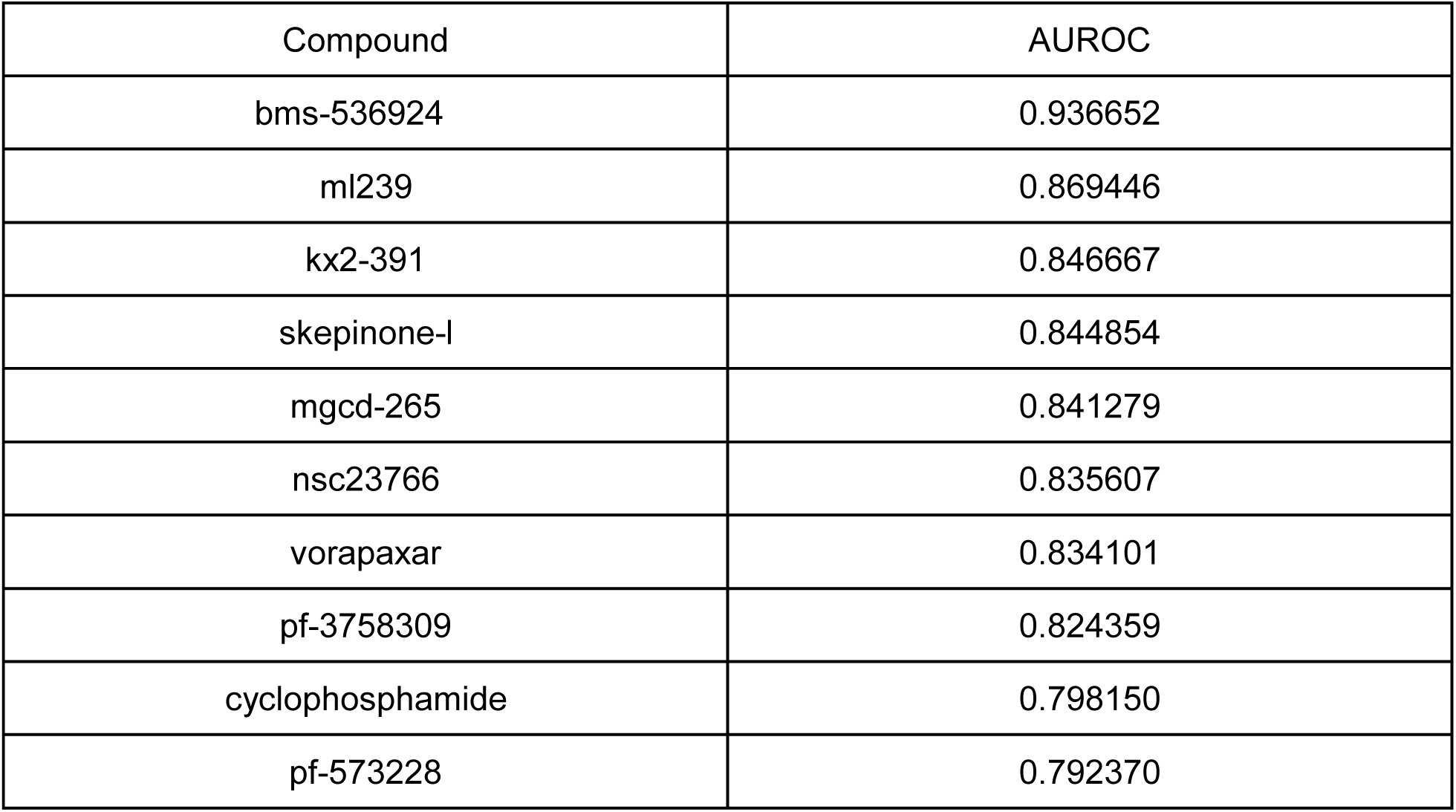
Compounds for which PACER predicted pathways with greatest precision. Evaluation was performed with known targets.

| Compound | AUROC |
| --- | --- |
| bms-536924 | 0.936652 |
| ml239 | 0.869446 |
| kx2-391 | 0.846667 |
| skepinone-l | 0.844854 |
| mgcd-265 | 0.841279 |
| nsc23766 | 0.835607 |
| vorapaxar | 0.834101 |
| pf-3758309 | 0.824359 |
| cyclophosphamide | 0.798150 |
| pf-573228 | 0.792370 |

**Table 2.** Top 10 pathways predicted by PACER for the compound ‘cyclophosphamide’

| Pathway | Contains known target? | P-value of pathway (baseline method) |
| --- | --- | --- |
| Fanconi anemia pathway | NO | 0.2670 |
| CXCR4-mediated signaling events | YES | 1 |
| TCR signaling in naive CD4+ T cells | YES | 1 |
| IL2 signaling events mediated by PI3K | YES | 1 |
| ATR signaling pathway | NO | 1 |
| TCR signaling in naive CD8+ T cells | YES | 1 |
| Class I PI3K signaling events | NO | 1 |
| p53 pathway | YES | 0.3185 |
| BARD1 signaling events | NO | 1 |
| EPO signaling pathway | NO | 1 |

**Table 3.** Top 10 pathways predicted by PACER for the compound ‘nsc23766’

| Pathway | Contains known target? | P-value of pathway (baseline method) |
| --- | --- | --- |
| Signaling events mediated by focal adhesion kinase | YES | 0.0001 |
| Integrins in angiogenesis | YES | 0.0716 |
| Alpha4 beta1 integrin signaling events | YES | 0.0084 |
| a4b7 Integrin signaling | NO | 0.1006 |
| Nectin adhesion pathway | YES | 0.0521 |
| Integrin-linked kinase signaling | YES | 0.4421 |
| Neurotrophic factor-mediated Trk receptor signaling | YES | 0.0080 |
| Integrin family cell surface interactions | NO | 0.0003 |
| Beta5 beta6 beta7 and beta8 integrin cell surface interactions | NO | 0.0013 |
| Plexin-D1 Signaling | NO | 0.0364 |

We note that the AUROC values reported here are likely to be underestimates, as there is literature evidence for some of the reported pathways being associated with the compound, even though the pathway does not include a known target (and is thus considered a false positive in our AUROC estimate). For example, our method identified the ‘Fanconi anemia pathway’ as being associated with compound ‘cyclophosphamide’ (**Table 2**). The Fanconi anemia pathway is one of the major DNA damage response pathways disrupted in breast cancer^38^, and is known to play an important role in cancer treatment by DNA crosslinking agents such as cyclophosphamide^39, 40^. PACER also identified ‘EPO signaling pathway’ to be associated with cyclophosphamide. Cyclophosphamide treatment has been reported to affect EPO receptor expression in murine erythropoiesis^41^. Although we found no existing study to corroborate the PACER-predicted association between cyclophosphamide and the ‘ATR signaling pathway’, the cyclophosphamide analogue, mafosfamide, has been reported to activate the ATM/ATR-Chk1/Chk2 pathway^42^. In addition, the predicted association of cyclophosphamide with ‘Class I PI3K signaling events’ is supported by reports of the compound activating the PI3K/Akt/mTOR signaling pathway in the ovary^43^.

The Rac1-specific inhibitor nsc23766 was also identified by PACER as being associated with several pathways that include a known target (**Table 3**), as well as some pathways that do not but whose association is supported by literature evidence. For example, the pathway ‘Beta5 beta6 beta7 and beta8 integrin cell surface interactions’ was predicted as being associated with this compound, and even though the latter's known target is not in this pathway, various lines of evidence support the association. A study of cholangiocarcinoma found beta6 integrin to promote invasiveness by activating Rac1, and that the compound nsc23766 is able to suppress this invasiveness^44^ and can be used to identify poor prognostic HER2 amplified breast cancer patients^45^. Beta8 integrin influences Rac1 levels to promote cell invasiveness in glioblastoma^46^. Also, Beta8 integrin is reported to activate Rac1 signaling in endometrial epithelial cells^47^. In addition, ncs23766 was predicted to be associated with ‘a4b7 Integrin signaling’. While this pathway does not directly include the compound's target, Rac1 was previously shown to induce a4b7-mediated T cell adhesion to MAdCAM-1^48^, supporting the predicted association. Furthermore, ‘Plexin-D1 Signaling pathway’ was found by PACER to be associated with ncs23766. Plexin-D1 is a receptor for SEMA4A which inhibits Rac activation^49^. Thus, the anecdotal observations made above suggest that compound-pathway predictions made by PACER may sometimes be worth pursuing even if the compound's target is not included in the pathway.

We further evaluated PACER based on the identified pathways for compounds with similar chemical structures. We performed this evaluation under the assumption that compounds with similar chemical structures tend to be associated with the same pathways. To this end, we calculated the Tanimoto similarity between each pair of compounds according to their SMILE specifications. We regarded two compounds as having similar chemical structures if their Tanimoto similarity is larger than 0.8. This threshold was used in previous work to indicate high Tanimoto similarity^50^. For a given compound, we used PACER to rank other compounds according to the Spearman correlation coefficient between their rankings of pathways, and asked if this ranking was predictive of chemical structure similarity. For each compound, an AUROC score was computed to measure this predictive ability. We repeated this evaluation with the three different methods of ranking pathways for a compound: PACER, the baseline method, and the method of Huang *et al.* **Figure 2F** shows the number of compounds for which the AUROC is above a specified threshold, for each of the three methods. We found that PACER achieves better performance in identifying compounds with similar chemical structure based on similarity of pathway ranking. For example, 21 (of 42) compounds yielded an AUROC greater than 0.8 when using PACER, compared to 10 compounds meeting the same criterion when using the baseline method. As compounds with similar chemical structure tend to be functionally similar, our results demonstrate that PACER can be used to identify similar compounds by integrating prior network information into chemosensitivity data.

We also compared the associations predicted by the three methods to those identified from an external data set. To this end, we mined the Library of Integrated Network-Based Cellular Signatures (LINCS) L1000 data^23^, which reports genes differentially expressed upon treatment of various cell lines with a compound. For each compound in our analysis that is also included as a perturbagen in the L1000 compendium, we established a LINCS-based benchmark of significantly associated pathways. This was based on a Fisher's exact test (*p*-value ≤ 0.05) between pathway genes and the most differentially expressed genes from treatments with the same compound (see Methods). We required this criterion to be met in at least one of the cell lines for which data was available from LINCS. We then assessed the concordance between this set of LINCS-based compound-pathway associations and those predicted by either method presented above. We recognize that this is not an ideal benchmark: LINCS data points to genes (and, indirectly, to pathways) that are differentially expressed in response to treatment, while PACER and the compared methods base their pathway predictions on genes that have basal expression levels across cell lines that correlate with chemical response. At the same time, we expect the pathways affected by chemical treatment to also be, to an extent, involved in interpersonal variation of chemosensitivity, making this a suitable evaluation procedure. This was inspired by similar observations in cancer biology: genes and pathways disrupted in cancer tissues overlap with genes and pathways whose mutation status in germline non-tumor samples is informative about disease susceptibility and progression.

To test whether the significant pathways identified from LINCS data agree with the pathways predicted by one of the methods being evaluated (based on chemical response variation in CCLE cell lines), we counted the compounds for which the two sets of predicted pathways overlapped significantly (Fisher's exact test *p*-value ≤ 0.05). As shown in **Figure 2G**, the PACER approach predicts pathways concordant with the corresponding LINCS-based benchmark for more compounds, compared to the baseline method and that of Huang *et al.*^16^ For instance, when the baseline method used an FDR threshold of 10% to designate significant pathway associations for each drug, and the PACER method predicted the same number of pathways, the latter's predictions were concordant with the LINCS-based benchmark for 110 of the 481 compounds, a nearly two-fold improvement over the baseline method's predictions. Our evaluations actually provide evidence for the above-mentioned possibility that pathways predictive of drug sensitivity overlap with genes that mediate drug response. In fact, we found 86 compounds for which the pathways identified from basal expression correlations and the pathways identified from LINCS signatures overlap with FDR < 5%.

After observing the substantial improvement of PACER, we then investigated whether the performance of PACER is stable when only using experimental derived protein-protein interactions as input. We found that the performance of PACER, as per the three evaluations presented above, was stable when only using experimental derived protein-protein interactions as input (**Suppl. Figure 4-6**). We further demonstrated that the result of our method is robust to different numbers of top response-correlated genes used in PACER, as shown in **Suppl. Figures 7-9**. We compared different values for *k* in the top *k* genes chosen by PACER. We found that for two of the three evaluation schemes, results were comparable when using *k*=100, 150, 200, 250 and 300. For the third evaluation scheme, results were comparable when using k=200, 250, and 300. This demonstrates the stability of the algorithm's performance to different but reasonable values of *k* in its choice of top *k* response-correlated genes.

## DISCUSSION AND CONCLUSION

We have shown that embedding prior knowledge in a gene network can more accurately identify compound-pathway. Our new method, called PACER, identified many compound-pathway associations that are supported by known compound targets as well as literature evidence. Due to its unique ability to incorporate any suitable compendium of gene interactions, our approach may provide complementary insights into drug mechanisms of action.

Historically, pathways associated with a particular gene set are identified by using popular statistical methods such as Gene Set Enrichment Analysis^51^, Fisher's exact test (DAVID^52^) or the Binomial test (Reactome^53^). These tools test the overlap between differentially expressed genes and pathway members. They may also be applied to the set of drug-response-correlated genes (RCGs) analyzed here. Ingenuity Pathway Analysis^54^ is another related tool, which utilizes information about causal interactions between pathway members. Our study is similar to the above tools in that PACER also seeks to find pathways implicated by a gene set. However, our approach differs from these existing tools in that known molecular interactions (e.g., PPI) among different genes are taken into consideration. Thus, a gene set, be it the RCGs of a compound or the members of a pathway, is not treated merely as the sum of its parts, but also includes the relationships among those parts. Since the dominant theme in existing approaches is assessment of overlaps between two gene sets (MSigDB, DAVID, and Reactome adopt variations on this theme), our extensive comparisons between PACER and the baseline method of Fisher's exact test shed light on the relative merits of the new approach. A related line of work aims to identify differentially expressed subnetworks in a given interaction network, e.g., KeyPathwayMiner^55^, but these studies are only superficially relevant to our work since we aim to prioritize existing pathways instead of finding new pathways.

We consider two potential reasons for the strong performance of PACER. First, it is widely appreciated that a chemical compound not only affects individual genes, but also combinations of genes in molecular networks corresponding to core processes, such as cell proliferation and apoptosis. Our method postulates that even if the RCGs and a pathway may only have a few genes in common, they may be close to each other in the network. Although current compound pathway maps are incomplete, much relevant information is available in public databases of human molecular networks. While traditional pathway enrichment analysis methods like Fisher's exact test identify pathways according to the number of shared genes, PACER prioritizes pathways based on their proximities to RCGs in molecular networks. Second, manually curated pathways may have arbitrary boundaries due to the need to capture knowledge at different levels of detail. Consequently, identifying drug related pathways might be hindered by pathway boundaries. By leveraging the prior knowledge in molecular networks, PACER is more robust to pathways with different boundaries, thus improving the sensitivity of detecting compound-pathway associations.

We see many opportunities to improve upon the basic concept of PACER in future work. First, although the current PACER framework was developed in an unsupervised fashion, the scores assigned to each pathway for the given gene set can be used as the feature and plugged into off-the-shelf machine learning classifiers for compound-pathway association identification.

Second, although this study focused on chemosensitivity response, the PACER method is broadly applicable to testing the association between two sets of genes according to their proximity in the network. Finally, although we use gene expression data as the molecular profile of each cell line, it might be interesting to test our method based on other molecular data such as somatic mutations and copy number alterations.

